# RAD54 is essential for RAD51-mediated repair of meiotic DSB in Arabidopsis

**DOI:** 10.1101/2020.06.09.142000

**Authors:** Miguel Hernandez Sanchez-Rebato, Alida M. Bouatta, Maria E. Gallego, Charles I. White, Olivier Da Ines

## Abstract

An essential component of the homologous recombination machinery in eukaryotes, the RAD54 protein is a member of the SWI2/SNF2 family of helicases with dsDNA-dependent ATPase, DNA translocase, DNA supercoiling and chromatin remodelling activities. It is a motor protein that translocates along dsDNA and performs multiple functions in homologous recombination. In particular, RAD54 is an essential cofactor for regulating RAD51 activity. It stabilizes the RAD51 nucleofilament, remodels nucleosomes, and stimulates homology search and strand invasion activity of RAD51. Accordingly, deletion of RAD54 has dramatic consequences on DNA damage repair in mitotic cells. In contrast, its role in meiotic recombination is less clear.

RAD54 is essential for meiotic recombination in Drosophila and *C. elegans*, but plays minor roles in yeast and mammals. We present here characterization of the roles of RAD54 in meiotic recombination in the model plant *Arabidopsis thaliana*. Absence of RAD54 has no detectable effect on meiotic recombination in otherwise wild-type plants but RAD54 becomes essential for meiotic DSB repair in absence of DMC1. In Arabidopsis, *dmc1* mutants have an achiasmate meiosis, in which RAD51 repairs meiotic DSBs. Absence of RAD54 in *dmc1* mutants leads to meiotic chromosomal fragmentation. The action of RAD54 in meiotic RAD51 activity is thus downstream of the role of RAD51 in supporting the activity of DMC1. Equivalent analyses show no effect on meiosis of combining *dmc1* with the mutants of the RAD51-mediators RAD51B, RAD51D and XRCC2.

RAD54 is thus required for repair of meiotic DSBs by RAD51 and the absence of meiotic phenotype in *rad54* plants is a consequence of RAD51 playing a RAD54-independent supporting role to DMC1 in meiotic recombination.

**Author Summary:** Homologous recombination is a universal pathway which repairs broken DNA molecules through the use of homologous DNA templates. It is both essential for maintenance of genome stability and for the generation of genetic diversity through sexual reproduction. A central step of the homologous recombination process is the search for and invasion of a homologous intact DNA sequence that will be used as template. This key step is catalysed by the RAD51 recombinase in somatic cells and RAD51 and DMC1 in meiotic cells, assisted by a number of associated factors. Among these, the chromatin-remodelling protein RAD54 is a required cofactor for RAD51 in mitotic cells. Understanding of its role during meiotic recombination however remains elusive. We show here that RAD54 is required for repair of meiotic double strand breaks by RAD51 in the plant *Arabidopsis thaliana*, and this function is downstream of the meiotic role of RAD51 in supporting the activity of DMC1. These results provide new insights into the regulation of the central step of homologous recombination in plants and very probably also other multicellular eukaryotes.

## Introduction

Homologous recombination (HR) is a universally conserved DNA repair mechanism essential for maintaining genomic integrity and ensuring genetic diversity [1, 2]. In somatic cells, HR is used to repair DNA breaks caused by environmental and endogenous factors and is critical in the recovery of stalled and collapsed replication forks. In meiotic cells of the majority of studied eukaryotes, HR is essential for accurate chromosome segregation during the first meiotic division, also generating genetic diversity among meiotic products [3, 4].

Homologous recombination is a DNA repair pathway that involves the use of a homologous template for restoration of the original sequence. It is initiated by DNA double-strand breaks (DSBs) and subsequent resection of the 5’-ended strands of the DSB, generating long 3’ single-stranded DNA (ssDNA) overhangs [5]. The ssDNA overhangs are further coated by replication protein A (RPA) protecting them from nucleases and removing secondary structures. In a subsequent step, RPA is displaced by the recombinase RAD51 in somatic cells, or RAD51 and DMC1 in meiotic cells, forming a right-handed helical nucleofilament on the exposed single-stranded DNA (ssDNA) flanking the DSB [6, 7]. This helical nucleofilament performs the homology search and catalyses the invasion of a homologous DNA template sequence by the 3’-ended DNA strands, which are then extended through DNA synthesis. The resulting joint recombination intermediate can be processed through several different pathways eventually leading to separation of the recombining DNA molecules and restoration of chromosome integrity [1, 2].

The nucleoprotein filament is the active protein machinery for DNA homology search and strand exchange during HR. In somatic cells, the nucleoprotein filament is formed by the RAD51 recombinase. The *in vivo* assembly and disassembly of the RAD51 nucleoprotein filament is a highly dynamic process, regulated via the coordinated actions of various positive and negative factors, and notably, the RAD51 mediators [8, 9]. These proteins, involved in the regulation of the formation, stability and activity of the RAD51 nucleofilament, include the RAD51 paralogues and the SHU complex that are known to be essential RAD51 positive regulators (for reviews see [8–11]. The RAD51 paralogues are important for homologous recombination and DNA repair in somatic cells [9, 12]. In contrast, clear understanding of their roles during meiosis remains elusive. Budding yeast has two RAD51 paralogues, Rad55 and Rad57, which form a heterodimer, and are essential for meiotic recombination [13–15] and 4 Shu proteins (Psy3, Csm2, Shu1 and Shu3) forming the Shu/PCSS complex that is also required for Rad51 filament assembly and meiotic recombination [16]. Vertebrates, like *Arabidopsis thaliana*, have five RAD51 paralogues (in addition to DMC1): RAD51B, RAD51C, RAD51D, XRCC2 and XRCC3 which form different complexes [8–11]. Vertebrate mutants for any of the RAD51 paralogues are embryonic lethal and this has hampered the study of their meiotic phenotypes. Nevertheless, a number of studies have demonstrated that RAD51C and XRCC3 are essential for meiotic recombination both in vertebrates and plants [17–27]. In contrast, the possible meiotic roles of RAD51B, RAD51D and XRCC2 are less clearly understood. These three genes are highly expressed in meiotic tissues in animals [28–30] and plants [31–33]. In humans, mutation in XRCC2 has been linked to meiotic arrest, azoospermia and infertility [34] and absence of RAD51B or RAD51D lead to meiotic defects in the moss *Physcomitrella patens* and rice, respectively [35–37]. The Arabidopsis *xrcc2* mutant and, to a lesser extent *rad51b*, have been associated with increased meiotic recombination rates, but all three mutants are fully fertile and present no detectable meiotic defects [24, 38–40]. Vertebrate genomes also encode two Shu-related proteins, SWS1-SWSAP1, which form a complex dispensable for mouse viability but essential for meiotic progression [41]. To date, Shu proteins have not been identified in plants.

RAD51 nucleofilament activity is further supported by the highly conserved RAD54 protein, which belongs to the SWI2/SNF2 DNA helicase family. It is a dsDNA-dependent ATPase that uses energy from ATP hydrolysis to translocate along dsDNA. It is thus a motor protein and performs multiple functions in homologous recombination. In particular, RAD54 is an essential cofactor stimulating RAD51 activity. It has been shown to stabilize the RAD51 nucleofilament, remodel nucleosomes, stimulate homology search and strand invasion activity of RAD51, dissociate bound RAD51 after completion of strand exchange and even to catalyse branch migration [42–44]. Accordingly, deletion of RAD54 has dramatic consequences on DNA damage repair in mitotic cells (For reviews see [42–44]).

The role of RAD54 in meiotic recombination is less clear. In Drosophila and C*. elegans*, which exclusively rely on RAD51 (not DMC1), RAD54 is essential for meiotic recombination [45–47]. Yet, in most eukaryotes, meiotic HR is mediated by RAD51 and the meiosis-specific DMC1 [6, 48]. Interestingly however, while RAD51 is essential for homology search and strand invasion in mitotic cells, it only plays an accessory role for DMC1 in meiosis [49, 50]. Thus, DMC1 is the active meiotic recombinase but requires the support of RAD51 to function [49, 50]. Accordingly, data from budding yeast have demonstrated that Rad51 activity is downregulated during meiosis to favour Dmc1 catalysing DNA strand-exchange using the homologous chromosome as a template [49, 51–54].

In yeast, down-regulation of Rad51 activity is mediated by the coordinated phosphorylation of Hed1 and the Rad51-cofactor Rad54 by the meiosis-specific kinase Mek1 [51–57]. Hed1 is a meiosis-specific protein that binds to Rad51, impeding access of Rad54 and thereby restricting activity of Rad51 nucleofilaments in meiosis [52, 55, 56, 58]. Phosphorylation of Rad54 by Mek1 also reduces its affinity for Rad51 [51, 59]. Thus, both pathways downregulate Rad51 through inhibition of Rad51-Rad54 complex formation and this in turns favour Dmc1-dependent inter-homologue recombination. In accordance with this down-regulation, Rad54 is also not essential for Dmc1 activity and plays a relatively minor role in meiotic recombination in budding yeast [60–66]. This is however due to the presence of a second, Dmc1-specific Rad54 homologue, Rdh54/Tid1 [62, 64–66]. Biochemical and genetic experiments have demonstrated that Rdh54 preferentially acts with Dmc1 to promote inter-homologue recombination whereas Rad54 preferentially stimulates Rad51-mediated strand invasion for sister chromatid repair of excess DSBs [60, 61, 64, 67, 68].

In mouse, two RAD54 homologues, RAD54 and RAD54B, have been identified. Both are required for somatic recombination but neither is essential for meiotic recombination as single and double mutant mice are fertile, although RAD54 may be needed for normal distribution of RAD51 on meiotic chromosomes [69, 70]. To date in plants, only one RAD54 orthologue has been characterized (Arabidopsis locus AT3G19210). As in yeast and mammals, Arabidopsis RAD54 is essential for RAD51-mediated recombination in somatic cells. Absence of RAD54 leads to DNA damage hypersensitivity, strong reduction in homologous recombination efficiency and defects in pairing of homologous loci following DSB formation [71–76]. However, beyond the fact that Arabidopsis *rad54* plants are fertile, a role for RAD54 in Arabidopsis meiotic recombination has not been assessed. Given its essential role in RAD51-nucleofilament activity and its expression in meiocytes [32, 33] we hypothesized that RAD54 may also play an important role in meiotic recombination in plants.

Here, we present a detailed analysis of RAD54 function in meiotic recombination in Arabidopsis. Our data show that absence of RAD54 has no detectable effect on meiotic recombination in otherwise wild-type plants, but that RAD54 becomes essential for meiotic DSB repair in absence of DMC1. In Arabidopsis *dmc1* mutants, RAD51 repairs meiotic DSBs but does not produce chiasmata and absence of RAD54 in *dmc1* mutants leads to massive chromosome fragmentation (a “*rad51*-like” phenotype). RAD51 immunolocalization confirms that meiotic RAD51 nucleofilaments are formed (but non-productive) in *dmc1 rad54* double mutants. Strikingly, similar analyses show no effect on meiosis of combining *dmc1* with the mutants of the RAD51-mediators RAD51B, RAD51D and XRCC2.

Altogether our data demonstrate that RAD54 is required for RAD51-dependent repair of meiotic DSBs in Arabidopsis in the absence of DMC1. The absence of detectable meiotic phenotype in *rad54* plants is thus a consequence of RAD51 playing only a supporting, non-catalytic role in meiotic recombination and this role is RAD54-independent. Our findings have several interesting implications for the regulation of the strand invasion step during meiotic recombination in Arabidopsis, which are further discussed.

## Results

### RAD54 is essential for somatic DNA repair

RAD54 is instrumental for homologous recombination in both mitotic and meiotic cells in many organisms (see above). In plants, previous analyses have also demonstrated a role of RAD54 in RAD51-mediated DSB repair in somatic cells, while the observation that *rad54-1* Arabidopsis mutant plants are fertile showed that the RAD54 protein does not play an essential role in Arabidopsis meiosis [71–76]. However, the existence of more subtle evidence for meiotic roles of RAD54 has not yet been assessed in plants. In addition to using the previously characterised *rad54-1* allele, we have characterised a second RAD54 T-DNA insertion allele (SALK_124992), which we have named *rad54-2* (Figure 1A). The exact genomic structure of the T-DNA insertion in the *rad54-2* allele was verified by PCR and sequencing (Figure 1A) and homozygous mutant lines were analysed by RT-PCR to confirm the absence of the respective transcripts (Figure 1B). In *rad54-2*, the T-DNA is inserted in exon 4 of the *RAD54* gene. This insertion is flanked by T-DNA LB sequences in opposite orientations and is associated with a deletion of 11 bp of the *RAD54* exon 4 sequence (Figure 1A). No transcript was detected with primers spanning the T-DNA insertion site, confirming the absence of full-length transcript (Figure 1B), although as commonly observed in the insertions, a transcript could be detected in *rad54-2* downstream of the T-DNA insertion. Sequence analysis showed that an in-frame stop codon is present in the upstream T-DNA left border, 24 bp after the chromosome-T-DNA junction (Figure 1A). Thus, a protein of the first 285 amino acids (out of 910) of RAD54 fused to 8 amino acids translated from the first 24 nt of the T-DNA LB could potentially be expressed from the *rad54-2* allele. If present, this protein would lack all of the described essential domains for RAD54 activity.

**Figure 1.**
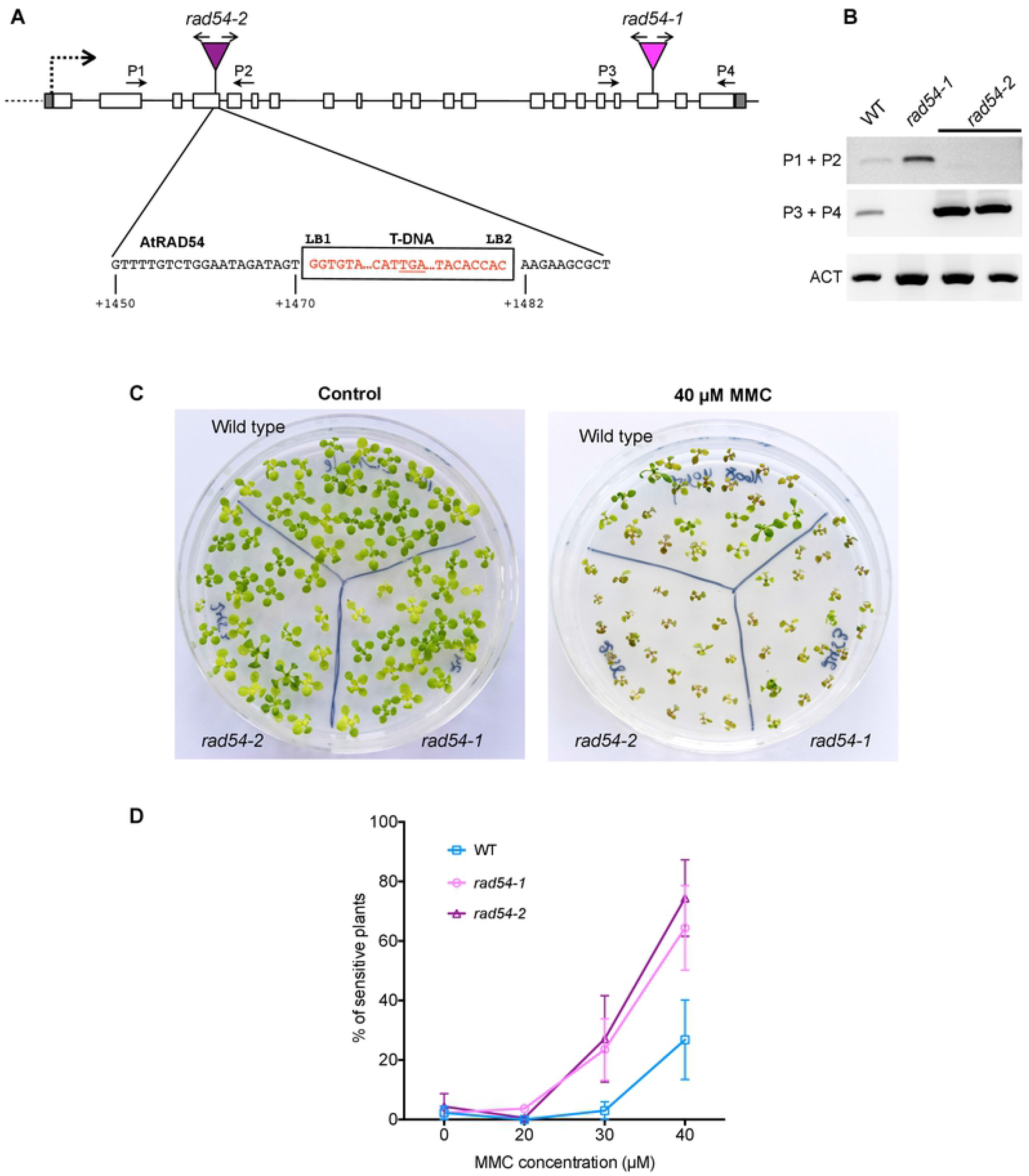
Characterisation of *rad54-2* T-DNA insertion mutant and sensitivity to MMC. (A) Structure of *AtRAD54* (At3g19210) and the *rad54-1* and *rad54-2* T-DNA insertion mutant alleles. Boxes show exons (unfilled) and 5’ and 3’UTRs (grey fill). The positions of the T-DNA insertions in the two alleles (inverted triangles) is indicated, with arrows above showing orientation of the left borders, and the sequences of the *rad54-2* T-DNA/chromosome junctions below. The *rad54-2* T-DNA insertion is flanked by two left borders (LB1, LB2) and accompanied by a 11 bp deletion in exon 4. An in-frame TGA STOP codon in *rad54-2* is underlined. Numbering under the sequences is relative to the *RAD54* start codon. (B) RT-PCR analyses of transcripts of *rad54-1* and *rad54-2*. Amplification of the actin transcript (ACT) was used as a control for RT-PCR. Positions and orientations of the PCR primers are shown on the diagrams. (C-D) Sensitivity of *rad54-1* and *rad54-2* plants to MMC. (C) Two-week-old seedlings grown without, or with 40 μM MMC are shown. (D) Sensitivity of the seedlings was scored after 2 weeks (see Materials and Methods) and the percentages of resistant plants (plants with more than 3 true leaves) are shown. Symbols are mean ± s.e.m of at least 3 independent experiments.

The *rad54-1* and *rad54-2* plants were used to confirm the role of RAD54 in DSB repair and homologous recombination in somatic cells by testing the sensitivity of the mutants to the DNA damaging agent Mitomycin C (MMC; Figure 1C-D). MMC is known to form DNA interstrand cross-link adducts, which produce DNA strand breaks *in vivo*. The importance of homologous recombination in the repair of DNA cross-links has led to the use of MMC hypersensitivity as a test for HR capacity in a number of organisms. In Arabidopsis, this is seen in the MMC hypersensitivity of many homologous recombination-deficient mutants [24, 26, 50, 74, 77]. As previously shown, *rad54-1* plants display clear hypersensitivity to MMC [74] (Figure 1C and D). MMC hypersensitivity is also seen in *rad54-2* plants, particularly visible at MMC doses of 30 and 40 μM (Figure 1C and D) and confirming the importance of RAD54 in homologous recombination in somatic cells.

### Absence of RAD54 does not affect meiotic progression

Meiotic defects are usually reflected in reduced fertility and thus in a reduction in seed number in Arabidopsis [78]. We thus monitored number of seeds per silique in our two *rad54* mutant lines and found, as expected, no fertility defects in either *rad54-1* or *rad54-2* (Figure S1). The mean seed number per silique was 56 seeds per silique for both *rad54-1* (n = 40 siliques) and *rad54-2* (n = 80), while wild-type siliques contained on average 58 seeds per silique (n = 40 for RAD54-1 and n = 60 for RAD54-2) (Figure S1). These small differences are not statistically significant (p > 0.05; unpaired, two-tailed Mann-Whitney test). In agreement with previous results [74], this confirms that RAD54 is not instrumental for meiosis in plants, notwithstanding its importance in somatic recombination. This conclusion was further supported through cytogenetic analyses of 4’,6-diamidino-2-phenylindole (DAPI) stained chromosomes through male meiosis. Wild-type Arabidopsis meiosis has been well described and the major stages are shown in Figure 2. During prophase I, meiotic chromosomes condense, pair, recombine and undergo synapsis. Full synapsis of homologues is seen at pachytene (Figure 2A). Chromosomes further condense and five bivalents (two homologous chromosomes attached by sister chromatid cohesion and chiasmata) are visible at metaphase I (Figure 2B). Each chromosome then separates from its homologue, leading to the formation of two groups of five chromosomes easily visualised at metaphase II (Figure 2C). Meiosis II proceeds and gives rise to 4 balanced haploid nuclei (Figure 2D). In *rad54* mutants, meiotic stages appear indistinguishable from the wild-type, resulting in the expected 4 haploid meiotic products (Figure 2E to L). Thus, meiotic progression is not affected by absence of RAD54.

**Figure 2.**
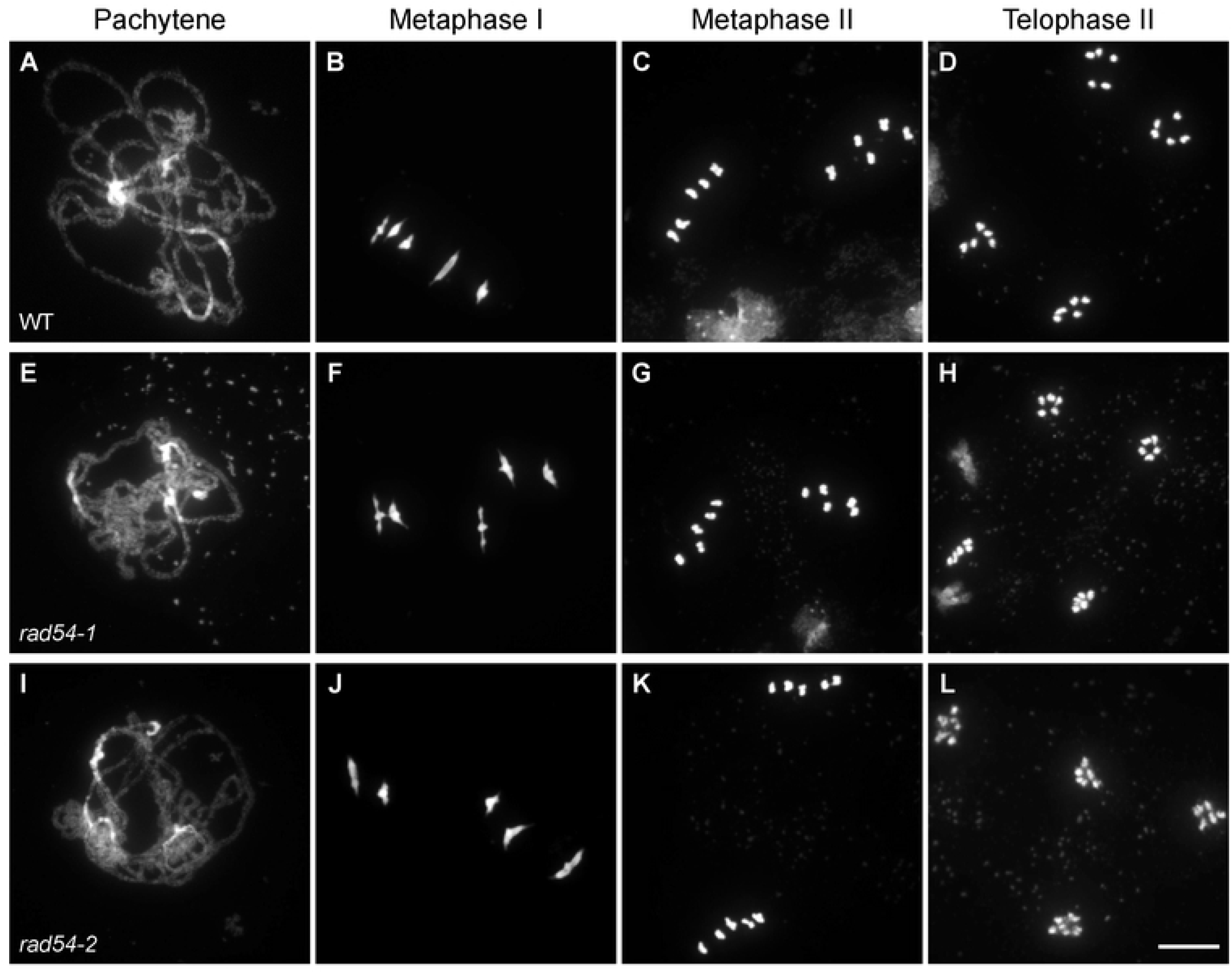
Both *rad54-1* and *rad54-2* mutants have WT meiosis. Chromosome spreads of male meiocytes in wild type (A-D), *rad54-1* (E-H) and *rad54-2* (I-L). Pachytene (A,E,I); Metaphase I (B,F,J); Metaphase II (C,G,K); Telophase II (D,H,L). Chromosomes were spread and stained with DAPI. (Scale bar = 10 μm).

### Absence of RAD54 does not affect crossover recombination rate and interference

We next sought to analyse more closely the impact of RAD54 on meiotic recombination by measuring meiotic CO rates in genetic intervals marked by transgenes encoding fluorescent marker proteins expressed in pollen (FTLs; [79, 80]). Combined with mutation of the *QUARTET1* gene (*qrt*) which prevents separation of the four pollen grains [81], these FTL lines permit direct measurement of recombination between the linked fluorescent markers by scoring tetrad pollen fluorescence [79, 80]. We determined CO rates in two adjacent intervals on chromosomes 1 (I1b and I1c) and 2 (I2f and I2g) in wild-type and *rad54-2* mutant plants. In wild-type plants, I1b (1.8 Mb) spans 10.3 cM and I1c (4.1 Mb) 22.2 cM (Figure 3 and Table S1). No difference in recombination frequency was observed for either interval in *rad54-2* mutants with 9 cM and 22.7 cM for I1b and I1c, respectively (Figure 3 and Table S1). Analyses of two additional intervals, I2f (0.7 Mb) and I2g (0.4 Mb), on chromosome 2 confirmed this result, with no significant difference in recombination frequency observed between the wild-type and *rad54-2* mutants (6.8 cM to 6.9 cM for I2f and 4.3 cM to 4.9 cM for I2g; Figure 3 and Table S1). We obtained similar results for *rad54-1* mutant plants with 6.5 cM and 4.7 cM in I2f and I2g, respectively (Supplemental Figure S2 and Table S1). In accordance with these results, we found a similar interference ratio (IR) in wild-type plants and *rad54* mutants for both intervals (IR I1bc : 0.35 in wild-type and 0.36 in *rad54-2;* IR I2fg : 0.9 in wild-type, 1 in *rad54-1* and 1 in *rad54-2;* p > 0.05, z-test).

**Figure 3.**
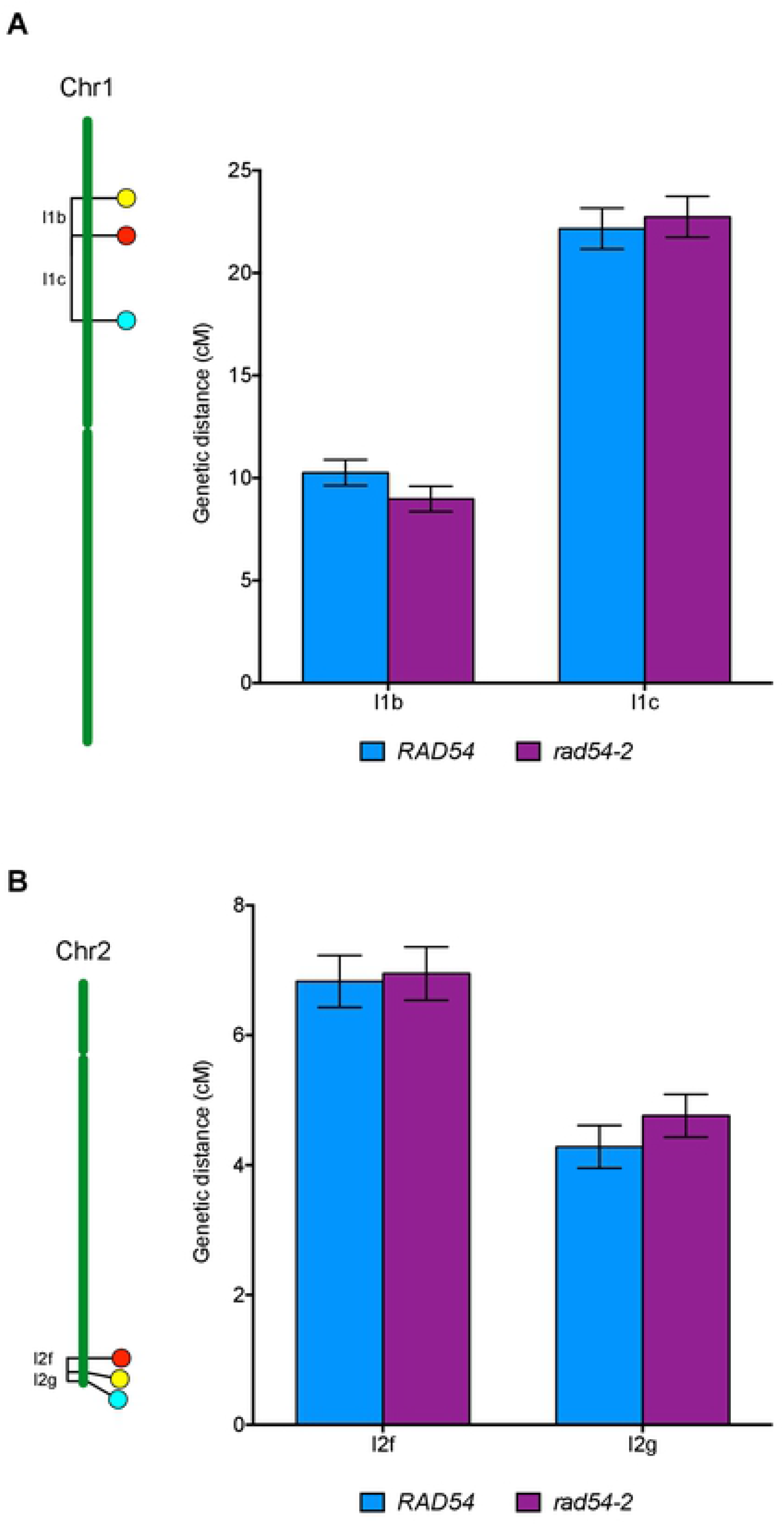
Crossing-over is not affected in *rad54-2* mutant meiosis. Genetic distances (in centiMorgans, cM) measured from fluorescent tetrad analyses in marked intervals on (A) chromosome 1 (I1b and I1c) and (B) chromosome 2 (I2f and I2g). Bars indicate mean ± SD. On all intervals, WT and *rad54* do not significantly differ (p<0.05; Z-test).

Thus, absence of RAD54 does not affect meiotic CO rates in at least 4 different intervals on 2 chromosomes. These results were further confirmed genome-wide through counting chiasmata in metaphase I of wild-type, *rad54-1* and *rad54-2* male meiocytes, which show means of 9.6 (SD = 1.3; n = 19), 9.6 (SD = 1.5; n = 25) and 9.1 (SD=1; n=19) chiasmata per meiosis, respectively (p > 0.05, unpaired two-tailed t-tests).

### RAD54 is essential for RAD51-dependent repair of meiotic DSB in absence of DMC1

These data confirm that RAD54 is not required for meiotic recombination in Arabidopsis, an *a priori* surprising conclusion given the importance of RAD54 in homologous recombination (see Introduction). Data from budding yeast have shown that RAD54 is not essential for meiotic recombination in presence of DMC1 and the DMC1-specific RAD54 homologue Rdh54 (Tid1) [51, 61, 64–66]. Instead, interaction of RAD54 with RAD51 is constrained during meiotic recombination in yeast and this represents a key point in the mechanisms leading to downregulation of RAD51 activity in meiosis [51, 58]. The RAD51-RAD54 pathway however becomes essential for sister chromatid repair in absence of DMC1 [60, 61]. We thus hypothesized that RAD54 may be essential for RAD51-mediated repair of meiotic DSB in Arabidopsis. To test this hypothesis, we analysed meiosis in the absence of DMC1. Meiosis in Arabidopsis *dmc1* mutants has been well described [82, 83] and the major stages are summarized in Figure 4. Absence of DMC1 leads to asynapsis and lack of CO. However intact univalents are observed in metaphase I owing to DSB repair by RAD51, most probably using sister chromatid (Figure 4E to H).

Analyses of *dmc1 rad54* double mutants show massive chromosome fragmentation in *dmc1 rad54* double mutants (Figure 4I to P), similar to that seen in *rad51* mutants (Figure 4Q to T). Thus, RAD51-dependent meiotic HR repair indeed depends upon the presence of RAD54 (Figure 4I to P). This effect is confirmed by the significant reduction of fertility caused by the absence of RAD54 in *dmc1* mutant plants (Figure S3). Accordingly, we propose that the absence of meiotic phenotype in *rad54* plants is a consequence of RAD51 playing only a RAD54-independent, non-catalytic role in supporting meiotic recombination by DMC1.

**Figure 4.**
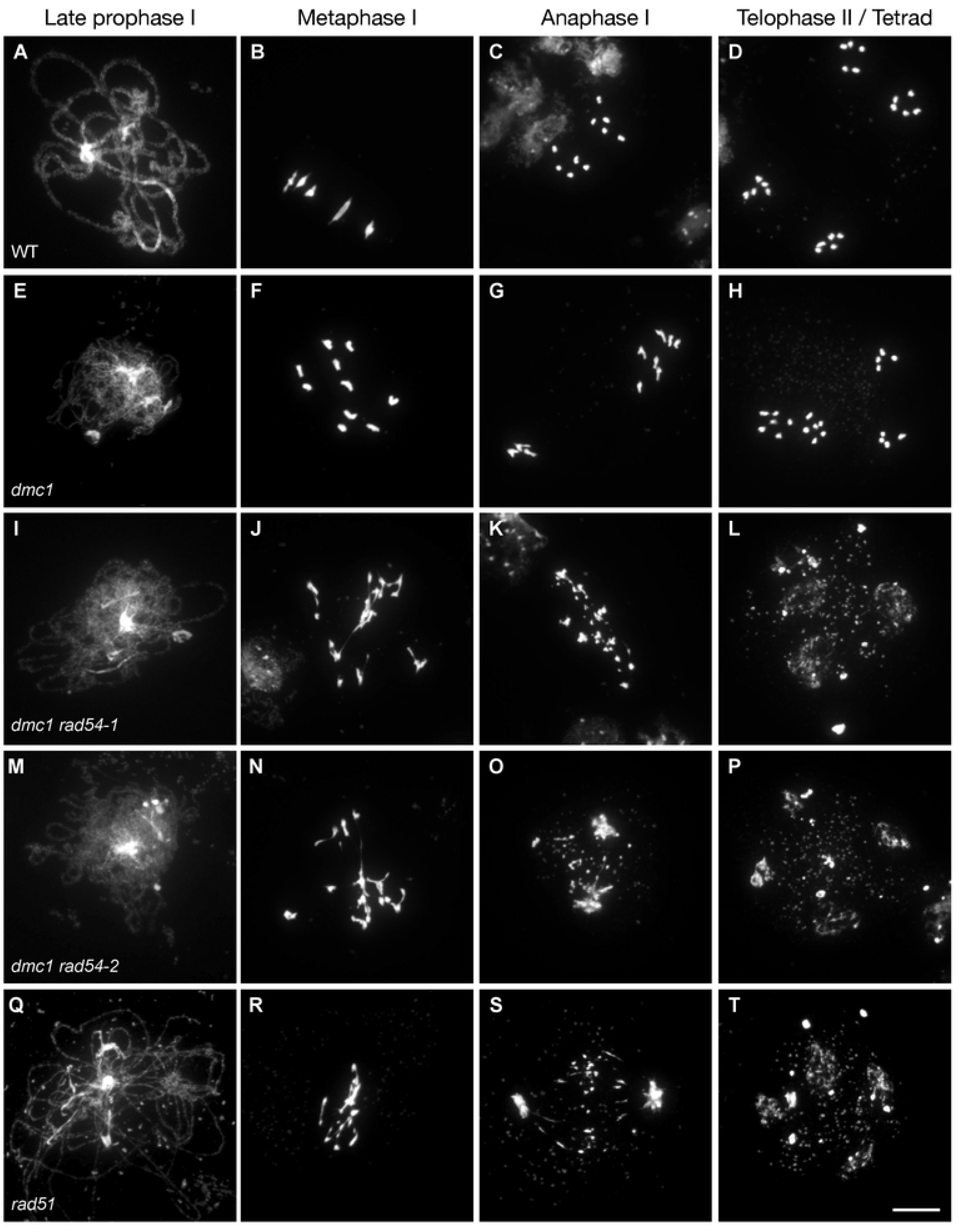
Absence of RAD54 leads to chromosome fragmentation in *dmc1* meiosis. Male meiosis is shown in (A-D) wild-type, (E-H) *dmc1*, (I-L) *dmc1 rad54-1*, (M-P) *dmc1 rad54-* 2, and *rad51* (Q-T). Chromosome spreads at late prophase I (A,E,I,M,Q), Metaphase I (B,F,J,N,R), Anaphase I (C,G,K,O,S) and Telophase II/Tetrad (D,H,L,P,T). Chromosomes were spread and stained with DAPI. (Scale bar = 10 μm).

### Absence of RAD54 blocks RAD51 catalytic activity rather than RAD51 focus formation

RAD54 is an essential cofactor for regulating RAD51 activity and has been implicated in both early and late steps of the HR pathway (see Introduction). Our data show that RAD54 is required for repair of meiotic DSB by RAD51 in Arabidopsis and the fact that RAD54 is not required in the presence of DMC1 suggests that the RAD54-dependence of RAD51 is downstream of that of the RAD51 nucleofilament in supporting DMC1 activity. To confirm the RAD54-independence of RAD51-nucleofilament formation in meiosis, we quantified meiotic RAD51 focus formation as a proxy for RAD51 nucleofilament formation in these plants. We performed co-immunolocalization of RAD51 and the axis protein, ASY1, in wild-type, *rad54*, *dmc1*, and *dmc1 rad54* meiocytes and counted the number of RAD51 foci throughout early prophase I (Figure 5). In wild-type meiocytes, we observed a mean of 91 ± 5 RAD51 foci (± S.E.M, n = 35). Similar numbers of RAD51 foci were observed in *rad54* (88 ± 6, n = 9) and *dmc1* (97 ± 3, n = 50) single mutant plants and importantly, the numbers of RAD51 foci were also unchanged in *dmc1 rad54* double mutants (94 ± 2, n = 56) (Figure 5). RAD51 nucleofilaments are still formed in the *dmc1 rad54* double mutants but are not productive. Hence, RAD54 acts downstream of RAD51 nucleofilament formation and its role must be in the activity of the nucleofilament, presumably in facilitating invasion of the donor DNA duplex [84]. We note also that this result concords with that fact that RAD54 is not needed for the essential role of the RAD51 nucleofilament in supporting DMC1 activity in meiotic HR.

**Figure 5.**
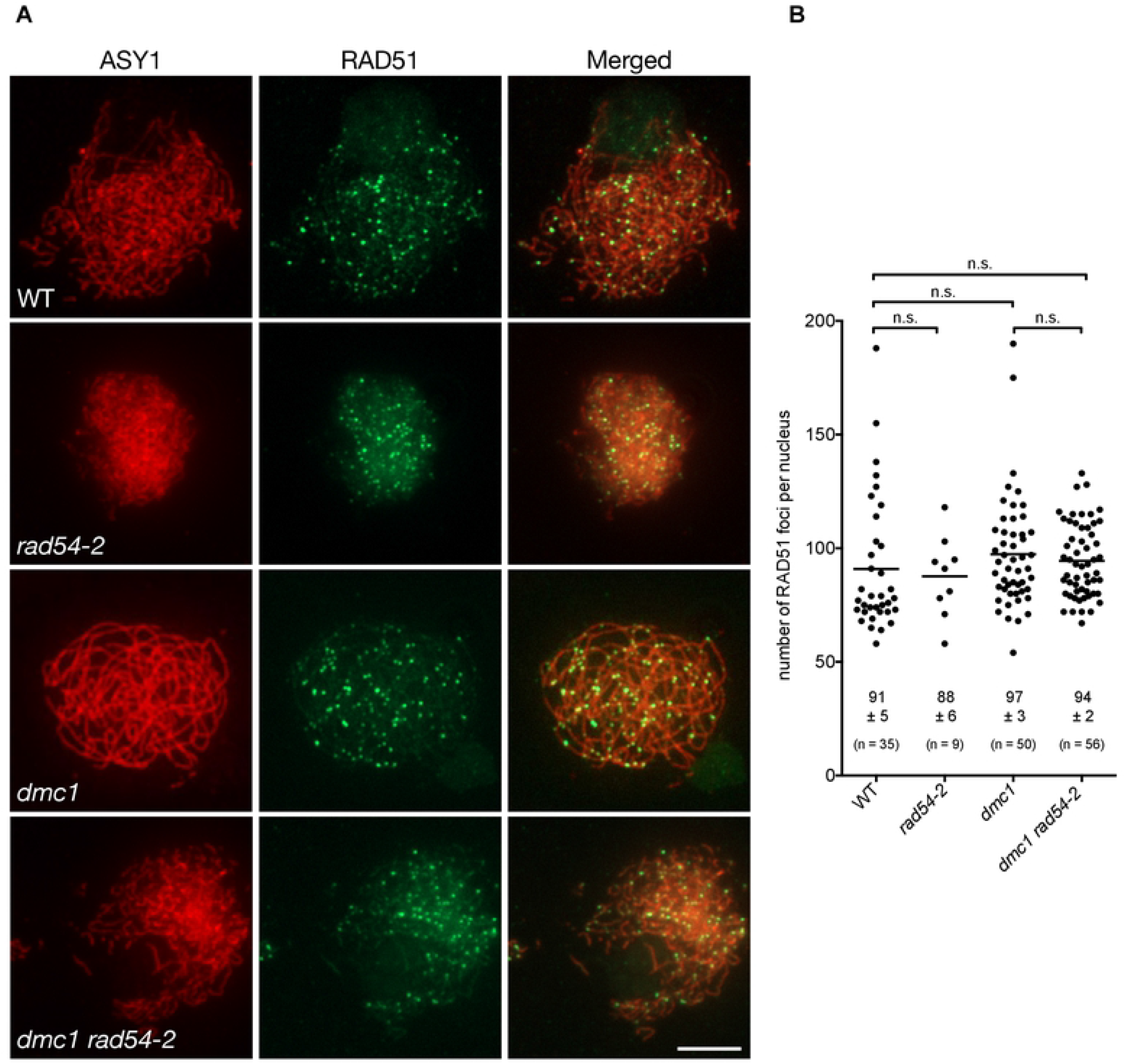
Absence of RAD54 does not affect numbers of meiotic RAD51 foci. (A) Co-immunolocalization of RAD51 (green) and the chromosome axis protein ASY1 (red) on leptotene/zygotene meiotic chromosome spreads. (Scale Bars: 5 μm). (B) Quantification of RAD51 foci per positive cell through early prophase I in wild-type, *rad54, dmc1*, and *dmc1 rad54-2* mutants. Means ± s.e.m are indicated. n.s.: not significantly different (p-value > 0.05, Kruskal-Wallis test).

### RAD51-dependent repair of meiotic DSB does not require RAD51 paralogues RAD51B, RAD51D and XRCC2

RAD51 nucleofilament activity is also extensively regulated by the RAD51 paralogues (see Introduction). In Arabidopsis, RAD51C and XRCC3 are essential for meiotic recombination, with absence of either leading to massive chromosome fragmentation [22–24, 26, 27]. In contrast, the roles of RAD51B, RAD51D and XRCC2 in meiosis are less clear and their absence does not lead to any obvious visible meiotic defects [24, 39, 40]. They are however expressed in meiotic tissues [31–33] and we have previously reported an increased meiotic recombination rate in Arabidopsis *xrcc2* (and to a lesser extent *rad51b*) mutants in two genetic intervals [39], suggesting potential roles for these paralogues during meiosis.

It thus appears possible, in analogy to RAD54 (above), that the absence of visible meiotic phenotype in *rad51b, rad51d* or *xrcc2* mutants could simply be a consequence of RAD51 strand-invasion activity not being required for meiotic recombination in the presence of DMC1.

We thus sought to test the impact of RAD51 paralogues in RAD51-dependent meiotic DSB repair by analysing meiotic progression in their absence in a *dmc1* mutant background (Figure 6). As described above, *dmc1* mutants are characterized by strong synaptic defects and lack of CO (Figure 6A-C). However, meiotic DSB are still repaired as seen in the presence of intact achiasmate univalents at metaphase I (Figure 6B), that segregate randomly at anaphase I (Figure 6C). These analyses did not show any detectable effects of the absence of RAD51B, RAD51D or XRCC2 in the *dmc1* mutant background (Figure 6D to L). In contrast, the expected chromosome fragmentation is observed in *xrcc3* mutant meiosis [26] and this is not affected by the additional absence of DMC1 (Figure 6M-O). Thus, despite being expressed in meiotic cells and playing key roles in RAD51 activity in somatic cells, RAD51B, RAD51D and XRCC2 are not required for RAD51-dependent meiotic DSB repair in Arabidopsis.

**Figure 6.**
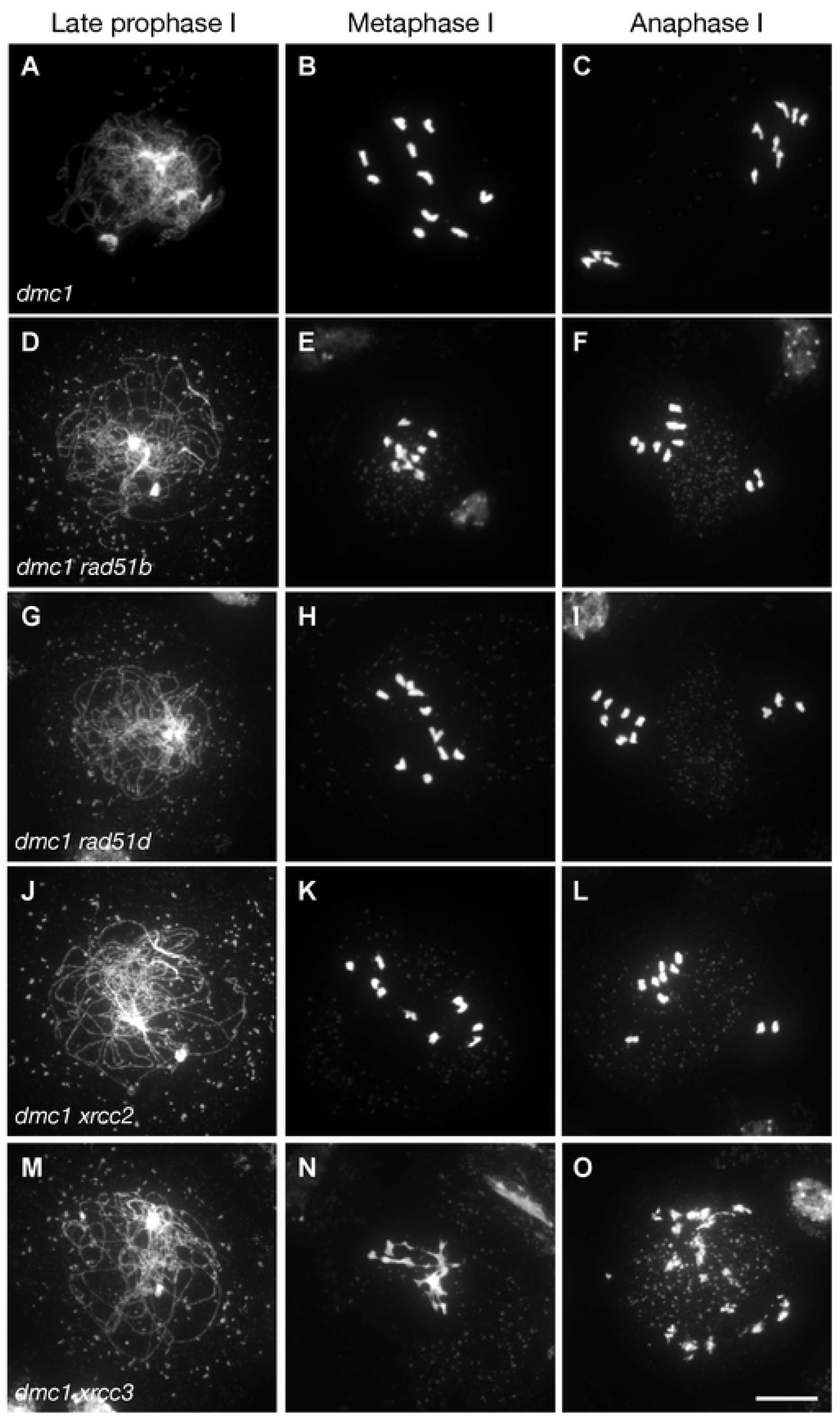
Absence of RAD51B, RAD51D or XRCC2 does not affect *dmc1* meiosis. Male meiosis is shown in (A-C) *dmc1*, (D-F) *dmc1 rad51b*, (G-I) *dmc1 rad51d*, (J-L) *dmc1 xrcc2*, and *dmc1 xrcc3* (M-O). Chromosome spreads at (A,D,G,J,M) late prophase I, (B,E,H,K,N) Metaphase I, (C,F,I,L,O) Anaphase I. Chromosomes were spread and stained with DAPI. (Scale bar = 10 μm).

## Discussion

Here, we provide evidence that Arabidopsis RAD54 is essential for meiotic double-strand break repair mediated by RAD51. This requirement for RAD54 is not observed in the presence of DMC1 as (all?) meiotic DSBs are repaired by DMC1 with RAD51 playing a supporting role to DMC1 in this process [49, 50, 59]. In the absence of DMC1 however, RAD51 catalyses the repair of meiotic DSB, leading to segregation of intact univalent chromosomes at meiotic anaphase I. Thus, absence of Arabidopsis RAD54 has no detectable effect on meiotic recombination in otherwise wild-type plants, but becomes essential for RAD51-dependent meiotic DSB repair in the absence of DMC1.

That this effect is not simply a reflection of a “mitotic” RAD51-dependent recombination context in *dmc1* meiosis is seen in the results of equivalent analyses with three RAD51 paralogue proteins, XRCC2, RAD51B and RAD51D, essential positive regulators of homologous recombination in somatic cells (reviewed in [9–11]). Mutants of these key RAD51-mediator proteins have no detectable meiotic phenotypes, beyond a mild meiotic hyper-rec phenotype reported for *xrcc2* and *rad51b* plants [24, 39, 40]. We report here that their absence does not visibly alter the meiotic phenotype of *dmc1* plants. Thus, in striking contrast to RAD54, the RAD51 paralogues RAD51B, RAD51D and XRCC2 are not required for RAD51-dependent meiotic DSB repair in Arabidopsis, despite being expressed in meiotic cells and playing key roles in somatic RAD51 activity.

RAD54 is a required cofactor for RAD51 activity and is thus instrumental for both mitotic and meiotic recombination in organisms lacking the meiosis-specific recombinase DMC1 [45–47]. The role of RAD54 in meiosis is however less clear in organisms expressing DMC1. Studies in budding and fission yeast have shown that Rad54 plays a relatively minor role in meiotic recombination [60–66]. This is however due to the presence of a second RAD54 homologue, Rdh54/Tid1. While both *rad54* and *rdh54* mutants form viable spores (albeit at reduced frequency), the *rad54 rdh54* double mutant rarely produces spores and is severely defective in meiotic recombination [62, 64–66]. These data reveal overlapping roles of Rad54 and Rdh54/Tid1 in meiotic recombination. In addition, Rdh54 preferentially acts with Dmc1 to promote inter-homologue recombination, whereas Rad54 preferentially stimulates Rad51-mediated strand invasion for sister chromatid repair [60, 61, 64, 68]. It is thus suggested that Rad54 is involved with Rad51 in sister chromatid repair of residual meiotic DSBs and this is in accordance with the recent demonstration of Rad51 being essential only to support Dmc1 and to repair residual DSBs after IH recombination is complete [49, 67, 85, 86].

In multicellular eukaryotes, evidence for a role of RAD54 homologues in meiosis however remains to be demonstrated. Mammals have two known RAD54 family members, RAD54 and RAD54B, neither of which appear to have important functions in meiosis, as mice lacking RAD54, RAD54B or both exhibit no, or only minor meiotic recombination defects [69, 70]. Our data demonstrate that RAD54 is essential for RAD51-mediated repair of meiotic DSBs in *dmc1* Arabidopsis. To our knowledge this is the first evidence of a clear meiotic role of RAD54 in a DMC1-expressing multicellular eukaryote. In Arabidopsis *dmc1* mutants, DSBs are repaired without formation of inter-homologue CO and this concords with the suggestion that RAD51 repairs meiotic DSB using the sister chromatid template [82, 83, 87]. Although this essential role is only observed in the absence of DMC1, we cannot exclude that the RAD51/RAD54 DSB repair pathway is also active (albeit weakly) in wild-type plants, possibly to repair excess DSBs as has been shown in yeast [60, 61]. Whether this pathway also exists in wild-type plants, remains however to be demonstrated.

Another conclusion inferred from our data is that Arabidopsis RAD54 is not necessary for DMC1 activity, either alone or as a RAD51 cofactor. That absence of RAD54 has no detectable effect on meiotic recombination in the presence of DMC1 tells us that RAD51’s function as an essential accessory factor for DMC1 is RAD54-independent. This conclusion concords with the reported absence of interaction between Arabidopsis RAD54 and DMC1 [74]. Yet, the DMC1 nucleofilament must perform homology search and strand invasion and this requires ATP-dependent DNA translocases (reviewed in [42, 44, 88]). We thus hypothesize that there exists a second, as yet unknown, DMC1-specific RAD54 homologue in plants. RAD54 is a SWI2/SNF2-remodelling factor that belongs to the SF2 helicase family, a number of which are encoded by the Arabidopsis genome [74, 76, 89], but to date only RAD54 (this work) has been found to play a role in meiosis.

Control of Rad51/Rad54 complex formation is used to downregulate Rad51 activity during meiosis in budding yeast, presumably to favour interhomolog recombination driven by Dmc1 [6, 49, 53, 54, 59]. This downregulation is largely achieved through preventing Rad51/Rad54 complex formation via two pathways involving two meiosis-specific proteins: the RAD51-binding protein Hed1 and the Mek1 kinase (which phosphorylates both RAD54 and Hed1) [51–53, 55, 56, 58]. Briefly, Mek1-mediated phosphorylation of RAD54 weakens RAD51-RAD54 interaction [51, 53] and binding of Hed1 to RAD51 also prevents association of RAD54 [52, 53, 55, 56, 58]. Interestingly, no apparent Hed1 or Mek1 orthologues have been identified in higher eukaryotes and in particular in plants. Several reports suggest that RAD51 is also down-regulated in Arabidopsis meiosis [50, 82, 83, 87, 90, 91], but the evidence for this remains indirect. Thus, whether RAD51 strand exchange activity is down-regulated during meiosis in higher organisms and if so, how this is achieved, is not clear. The absence of meiotic phenotype of Arabidopsis *rad54* mutants, together with the demonstration of the RAD54-dependence of meiotic RAD51 activity (in the absence of DMC1), supports the idea of a hypothetical RAD54-dependent control of RAD51 activity through modulation of the RAD54/RAD51 interaction. It also, however, invites speculation concerning whether it is necessary to invoke such a downregulation to explain numbers of CO vs non-CO recombination events in plants, and very likely in vertebrates. Previous work has shown that DMC1 is capable of catalysing repair of all meiotic DSB in Arabidopsis in strand-invasion mutants of RAD51 [50, 90], or as shown here, by blocking RAD51 activity through the absence of RAD54. In both of these contexts, no evidence of alteration of numbers nor distribution of meiotic recombination has been found.

In conclusion, we present here an essential role for RAD54 in supporting meiotic RAD51-mediated DSB repair in the absence of DMC1 in Arabidopsis. In striking contrast, testing of three other key RAD51 mediator mutants (*rad51b, rad51d, xrcc2*) did not reveal any detectable impact on *dmc1* meiosis, notwithstanding the fact that they are, like RAD54, needed for RAD51-dependent recombination in somatic cells. This RAD54-dependent, RAD51-mediated meiotic DSB repair is thus not the reflection of a simple “mitotic-like” RAD51 DSB repair in meiocytes lacking DMC1, but points to RAD54 acting downstream of the role of the RAD51 nucleofilament in supporting meiotic DMC1-mediated recombination. It will be of particular interest to further study in which context this pathway is activated in wild-type meiosis and also whether a similar pathway exists in other organisms outside the fungal taxa. Although further studies are needed to confirm whether (and how) RAD51 strand-invasion activity is downregulated during meiosis in plants, we speculate that this could be achieved through prevention of RAD54/RAD51 interaction, and/or via helicases dissociating precocious strand-invasion between sister chromatids, as has recently been shown in budding yeast [92].

## Materials and Methods

### Plant Material and Growth Conditions

All *Arabidopsis thaliana* plants used in this study were in the Columbia background. Seeds of the *rad54-2* (SALK_124992) [93] T-DNA insertion mutant were obtained through the Nottingham Arabidopsis Stock Centre and characterised in this study. For other mutants, we used the following alleles: *rad54-1* [74], *dmc1-2* [83], *rad51-1* [94], *rad51b-1 [24], rad51d-3* [39] and *xrcc2-1* [24]. Fluorescent-Tagged lines (FTLs) were: I1bc (FTL567-YFP/FTL1262-DsRed2/FTL992-AmCyan/*qrt1-2*), and I2fg (FTL800-DsRed2/FTL3411-YFP/FTL3263-*AmCyan/qrt1-2*) [79].

Seeds were stratified in water at 4°C for 2 days and grown on soil in a growth chamber. For *in vitro* culture, seeds were surface sterilised for 5 min with 75% Ethanol, 0.05% SDS, rinsed with 95% Ethanol for 5 min and air-dried. Sterilised seeds were then sown on half-strength Murashige and Skoog (MS) medium, stratified at 4°C for 2 days and placed in a growth cabinet. All plants were grown under 16h light /8 h dark cycles at 23°C and 60% relative humidity.

### Molecular Characterization of *rad54-2* T-DNA Insertion Mutants

The *rad54-2* (SALK_124992) mutant was genotyped using primers P1 and P2 to detect the wild-type loci and primers P1, P2, and Lba1 (SALK T-DNA Left Border specific primer) were used to detect the T-DNA insertion allele. The junctions of the T-DNA insertion in the *RAD54* locus (AT3G19210) were amplified by PCR and verified by DNA sequencing.

For semi-quantitative RT–PCR, total RNA was extracted from young buds of wild-type, *rad54-1* and *rad54-2* plants using RNeasy Plant mini Kit (QIAGEN), following the manufacturer’s instructions. 2 μg RNA were treated with RQ1 RNase-free DNase (Promega) followed by reverse transcription using M-MLV Reverse Transcriptase (Promega) according to the manufacturer’s instructions. PCR amplifications were eventually performed in homozygous lines showing the absence of full-length *RAD54* transcripts (Figure 1).

Sequences of primers used for genotyping and RT-PCR are listed in Supplemental Table S1.

### Mitomycin C Sensitivity Assays

For the MMC sensitivity assay, seeds were surface-sterilised and sown onto solid medium (half strength Murashige and Skoog salts, 1% sucrose, 0.8% agar) supplemented with 0, 20, 30 or 40μM Mitomycin C (SIGMA). Seeds were stratified in the dark for 2 days at 4°C, transferred to a growth cabinet and grown for two weeks. Sensitivity was then analysed in two-week-old seedlings by counting the number of true leaves as previously described [26]. Plants with more than three true leaves were considered as resistant. In each case, the number of leaves was counted on at least 25 seedlings in three to five independent experiments.

### Recombination measurement using Fluorescent-Tagged Lines (FTL) tetrad analysis

We used Fluorescent Tagged Lines to estimate male meiotic recombination rates at two pairs of genetic intervals: I1bc on chromosome 1 and I2fg on chromosome 2. For each experiment, heterozygous plants for the linked fluorescent markers were generated and siblings from the same segregating progeny were used to compare the recombination frequency between different genotypes. Slides and fluorescent tetrad analysis were performed as described by Berchowitz and Copenhaver [79]. Tetrads were counted and attributed to specific classes (A to L). Genetic distances of each interval were calculated using Perkins equation as follows: *X* = 100[(1/2Tetratype + 3Non-Parental Ditype)/*n*] in cM.

The Interference Ratio (IR) was calculated as described previously [79]. Briefly, for two adjacent intervals I1 and I2, two populations of tetrads are considered: those with at least one CO in I2 and those without any CO in I2. Genetic distance of I1 is then calculated for these two populations using the Perkins equation, i.e. *X1* (I1 with CO in I2) and *X*2 (I1 without a CO in I2). The Interference Ratio is thus defined as IR = *X*1/*X*2. An IR ratio <1 reveals the presence of interference while an IR ratio close to 1 reveals absence of interference. The Stahl Lab Online Tools was used for statistical analyses of the data.

### Arabidopsis male meiotic chromosome spreads

Meiotic chromosome spreads were prepared according to [95]. Whole inflorescences were fixed in ice-cold ethanol/glacial acetic acid (3:1) and stored at −20°C until further use. Immature flower buds of appropriate size were selected under a binocular microscope and incubated for 75-90 min on a slide in 100μl of enzyme mixture (0.3% w/v cellulase (Sigma), 0.3% w/v pectolyase (Sigma) and 0.3% cytohelicase (Sigma)) in a moist chamber at 37°C. Each bud was then softened for 1 minute in 20 μl 60% acetic acid on a microscope slide at 45°C, fixed with ice-cold ethanol/glacial acetic acid (3:1) and air dried. Slide were mounted in Vectashield mounting medium with DAPI (1.5 μg.ml^−1^; Vector Laboratories Inc.).

### RAD51 Immunolocalization in meiocytes

Spreads of PMCs for immunolocalization of RAD51 were performed as described previously [96]. Primary antibodies used for immunostaining were: anti-ASY1 raised in guinea Pig (1:500) [97] and anti-RAD51 raised in rat (1:500) [98]. Secondary antibody: anti-rat Alexa fluor 488; anti-rat Cy3 were used at 1:100 dilution.

### Microscopy

All observations were made with a motorised Zeiss AxioImager.Z1 epifluorescence microscope (Carl Zeiss AG, Germany) driven by the ZEN Pro software (Carl Zeiss AG, Germany). Photographs were taken with an AxioC.am Mrm camera (Carl Zeiss AG, Germany) and Zeiss filter sets adapted for the fluorochromes used. Image stacks were captured in three dimensions (x, y, z) and further processed and adjusted for brightness and contrast on ZEN Pro and ImageJ/FIJI software. RAD51 foci were counted on collapsed z-stack projections by using counting tool of the ZEN Pro software.

## Acknowledgements

We thank Gregory Copenhaver and Ian Henderson for FTL lines, Chris Franklin and Peter Schlögelhofer for providing the ASY1 and RAD51 antibodies, respectively. We thank members of the recombination group for their help and discussions.

## Supporting information

**Supplemental Figure 1. Fertility of *rad54-1* and *rad54-2* mutants.**

**(A)** pictures of wild-type and *rad54* mutant siliques. **(B)** Number of seeds per silique in Wild-type, *rad54-1* and *rad54-2* mutants. Each point represents the number of seeds in one silique. Bars indicate mean ± SEM. n.s. : not significantly different. P > 0.05 (unpaired, two-tailed Mann-Whitney test).

**Supplemental Figure 2. Genetic recombination in wild-type, *rad54-1* and *rad54-2* mutants measured using I2fg fluorescent-tagged lines.**

Genetic distances (in centiMorgans, cM) calculated from tetrad analysis of the I2f and I2g intervals on chromosome 2. Bars indicate mean ± SD. For both intervals, WT and *rad54* plants do not significantly differ (p<0.05; Z-test).

**Supplemental Figure 3. Fertility of *dmc1 rad54-1* and *dmc1 rad54-2* mutant plants.**

Number of seeds per silique in Wild-type, *dmc1, dmc1 rad54-1* and *dmc1 rad54-2* mutants. Each spot represents the number of seeds in one silique. Bars indicate mean ± SEM. ****: significantly different. P < 0.0001 (unpaired, two-tailed Mann-Whitney test).

**Table S1. FTLs raw data.**

Tetrad count for all tetrad categories for I1bc and I2fg intervals. Tetrad categories (a to l) were classified as described previously by Berchowitz and Copenhaver (2008).

